# Spatio-temporal protein interaction analysis using bimolecular fluorescence complementation in *C. elegans*

**DOI:** 10.1101/2025.02.28.640866

**Authors:** Charlotte J. Martin, John A. Calarco

**Affiliations:** Department of Cell and Systems Biology, University of Toronto, 25 Harbord Street, Toronto, Canada, M5S 3G5

## Abstract

Dynamic protein-protein interactions (PPIs) shape all aspects of cellular biology. Thus, significant efforts have been made to develop assays testing binary PPIs. The transparency of *C. elegans* makes it a great model organism for fluorescence-based PPI detection *in vivo*. However, to date, there is currently a lack of quantitative PPI assays that also provide information on the subcellular location of protein interactions in *C. elegans.* Here, we have made several modifications to the original bimolecular fluorescence complementation (BiFC) assay used in *C. elegans* to make it more quantitative and spatio-temporally controlled. First, transgenes are expressed at single copy, reducing the variability associated with multi-copy expression. Second, we have added bicistronic reference fluorescent proteins to each transgene, allowing for the normalization and quantification of the PPI. Finally, we have incorporated the auxin-inducible degradation system, allowing for small-molecule inducible control of the PPI signal. We demonstrate the utility of our modified BiFC assay by testing several model PPIs. Thus, we anticipate that our updated BiFC approach will expand the available tools for studying PPIs in *C. elegans*, but similar logic could be applied to other model organisms amenable to transgenesis and *in vivo* fluorescent imaging.

**Article Summary:** Protein-protein interactions (PPIs) play a central role in all facets of cellular biology. Here, we developed an improved assay to study PPIs in *C. elegans*, based on bimolecular fluorescence complementation (BiFC), where two halves of split-YFP can be reconstituted in an interaction-dependent manner. Our modifications include making the readout of the assay less variable and more quantitative, while also enabling signal to accumulate in an inducible manner. We envision that our updated BiFC approach will serve as a useful tool for *C. elegans* researchers interested in characterizing PPIs of interest *in vivo*.

## Introduction

The identification and characterization of protein-protein interactions is a major driver of discovery of the complex machinery of cells. Given their importance and centrality in cell biology, numerous approaches have been developed to identify protein interactions. For example, affinity- or immuno-purification of proteins of interest or proximity labeling approaches such as bioID can, when coupled with mass spectrometry, lead to the unbiased detection and identification of protein complexes (Dunham et al., 2012; Roux et al., 2012). Indeed, these approaches, when paired with functional genomics analysis and network visualization, can lead to insights and hypotheses about protein complexes and biological functions.

Complementary to these global protein-protein interaction approaches, more focused assays testing binary interactions have been developed for characterization of protein complexes. For example, proteins found in complexes are routinely identified by the immunoprecipitation of a protein and then subsequent identification of a specific interacting protein through western blotting (Burnette, 1981; Jeong et al., 2018; Towbin et al., 1979). A more high-throughput approach is the yeast two-hybrid assay, in which a DNA binding domain is fused to a bait protein and then a transcription activator domain is fused to a prey protein (Fields & Song, 1989). If the bait and prey protein interact the DNA binding domain and transcription activator domain are reconstituted, leading to the activation of a reporter gene in the yeast (Fields & Song, 1989). An additional high-throughput approach to assay binary interactions has been developed in mammalian cells called LUminescence-based Mammalian IntERactome (LUMIER) (Barrios-Rodiles et al., 2005). In this assay, the bait protein is fused to a *Renilla* luciferase enzyme (RL) and then the prey protein is immunoprecipitated via a fused Flag tag and the interaction of bait and prey protein is identified by the presence of light from RL enzymatic activity (Barrios-Rodiles et al., 2005).

In the nematode *Caenorhabditis elegans*, a number of these above-mentioned approaches have been utilized. To identify unknown interactors for a given protein, a popular method is affinity purification paired with mass spectrometry (AP-MS) (Remmelzwaal & Boxem, 2019). More recently, the introduction of proximity labelling paired with mass spectrometry through the biotin ligase TurboID has increased in utility and feasibility, even enabling the identification of protein interactions in specific cell types and subcellular locations (Artan et al., 2021; Branon et al., 2018; Holzer et al., 2022; Sanchez et al., 2021; Zhao et al., 2024). For binary interactions, the *C. elegans* interactome has been well studied through yeast two-hybrid assays (Y2H) (Simonis et al., 2009). However, Y2H data provides no information on the localization of an interaction and requires additional modifications to study certain proteins (e.g. membrane proteins). Moreover, given interactions identified via Y2H are not in the native organism, any detected interactions would need to be confirmed *in vivo* in *C. elegans*.

Due to its transparency, *C. elegans* is also a tractable model organism for use of fluorescence as a readout to visualize protein expression patterns, activation and subcellular localization. Moreover, fluorescent sensors of calcium and other metabolites have been developed and used to great effect in the worm (Banerjee et al., 2015; Ding et al., 2023; Tian et al., 2009; Woldemariam et al., 2019) Finally, fluorescence-based assays have been adapted for use in *C. elegans* to detect or validate protein-protein interactions *in vivo*. Recently, the Light-induced co-clustering technique was adopted in *C. elegans* (Kroll et al., 2021). CeLINC uses a CRY2/CIB1 light-dependent oligomerization system to cluster fluorescently tagged interacting proteins together, allowing for the visualization of binary protein-protein interactions *in vivo* (Kroll et al., 2021). CeLINC was effective in detecting known protein-protein interactions between cell polarity components (Kroll et al., 2021). The CeLINC system offers a number of advantages over assays such as Y2H. Specifically, interactions can be detected in their appropriate organismal context and in physiologically relevant cell types. However, one trade-off in CeLINC is that due to a requirement of recruitment to sites of oligomerization, interacting proteins are detected at locations distinct from where the proteins are normally localized (Kroll et al., 2021). Thus, spatial information about where protein interactions may be occurring is typically lost, as well as any influence from the subcellular compartment on the strength of an interaction.

Another fluorescent protein-based assay to detect protein-protein interactions that has been adapted for use in *C. elegans* is bimolecular fluorescence complementation (BiFC). BiFC is technique in which two proteins of interest are tagged with one of two halves of a split fluorescent protein (Hiatt et al., 2008; Hu et al., 2002; Shyu et al., 2008). If the two proteins interact, the two halves of the fluorescent protein are brought together, reconstituting a fluorescent signal (Hiatt et al., 2008; Hu et al., 2002; Shyu et al., 2008). When no interaction occurs, fluorescence should be absent or significantly diminished. Much like CeLINC, an advantage of the BiFC approach is that it allows the detection of binary PPIs directly in *C. elegans* and in physiologically relevant tissues. However, a key advantage over CeLINC is that the subcellular location where the query proteins are interacting is preserved. Despite its potential, there are several disadvantages of the BiFC system in its current state. First, the split YFP protein reconstitution is irreversible (Hiatt et al., 2008; Shyu et al., 2008). Therefore, if given enough time, the two halves of the split-YFP are capable of finding each other at some frequency independent of any protein-protein interactions, ultimately leading to a saturated signal for interacting proteins and the potential for increased background signal (Hiatt et al., 2008; Shyu et al., 2008). Second, in *C. elegans*, the method has only been implemented using multi-copy expression of the BiFC vectors (Bhan et al., 2020; Kalis et al., 2022; Tien et al., 2011; Van De Walle et al., 2019). This multi-copy expression can lead to variation in the fluorescent signal between different animals and even in different cells within the same animal due to differences in copy-number from extrachromosomal arrays (Hiatt et al., 2008; Shyu et al., 2008). These copy number differences and irreversible accumulation of split-YFP signal make quantification of a PPI challenging, and thus most BiFC readouts are typically qualitative in nature (Chen et al., 2007; Mathews et al., 2012; Niu et al., 2017; Pandey et al., 2017).

In this study, we present an updated version of the original BiFC assay in an effort to make it more user-friendly and to address some of its caveats. First, we have generated a series of BiFC expression vectors compatible with single-copy transgenesis thus reducing copy-number variation issues associated with the approach (Nonet, 2020). Second, using 2A peptides (Ahier & Jarriault, 2014), we co-express additional fluorescent proteins that are used to normalize protein expression in individual cells, facilitating a quantitative measure of the strength of the queried PPI. Finally, we have made our expression vectors compatible with the auxin-inducible degron system (Ashley et al., 2021; Zhang et al., 2015), enabling spatio-temporal control of the signal for the PPI being queried. We demonstrate the importance of these updates and the utility of our revised expression vectors by testing several model PPIs in neurons and muscle cells. Taken together, we anticipate that our modifications to the BiFC system will further facilitate its use in *C. elegans* research and expand the toolkit to query PPIs in this key model organism.

## Materials and Methods

### Strains and maintenance

Animals were grown using standard worm maintenance procedures at 21°C on NGM plates seeded with OP50-1 *E. coli* bacteria (Brenner, 1974) unless otherwise stated. For a complete list of strains used see **Table S1**.

### BiFC expression vector cloning

Plasmids were constructed using a variety of approaches. Initial vectors pCM4 and pCM5 containing worm codon optimized split-YFP fragment, mTagBFP2, mCherry, AID and 2A peptide coding sequences were ordered from GenScript (see Fig. 1 for schematic). The *rgef-1* (pan-neuronal) and *myo-3* (body-wall muscle) promoters were amplified from genomic DNA and the pCFJ104 plasmid (*pmyo-3::mCherry*), respectively, using primers oCM58-61. Plasmids pCM4 and pCM5 were digested with the restriction enzyme NheI and promoters were inserted via Gibson Assembly, creating plasmids pCM7-pCM10. The cDNAs encoding the basic region-leucine zipper (bZip) domains from mouse JUN and FOS (bJUN and bFOS) and a FOS bZip domain lacking a leucing zipper motif (bFOSΔZIP) were ordered as gBlocks from IDT (oCM87-89). Plasmids pCM7-pCM10 were digested with the restriction enzyme NotI. The bFOS and bJUN cDNAs were then inserted through Gibson assembly, creating plasmids pCM22-27. Then the plasmids were subcloned into the FLP recombinase mediated cassette exchange (RMCE) integration vector pLF3FShc (ordered from addgene) (Nonet, 2020). pLF3FShc was digested with the restriction enzyme SmaI and full expression transgenes were amplified from pCM22-27 by PCR using primers oCM136-38 and oCM141 and inserted via Gibson assembly, generating RMCE BiFC expression plasmids pCM33, pCM35 and pCM38-42. For insertion of *rgef-1* promoter into our RMCE BiFC expression vectors, we removed the *myo-3* promoter from pCM39 and pCM41-42 by digestions with restriction enzyme NheI and subcloned the *rgef-1* promoter either using T4 DNA ligase or Gibson Assembly (amplified *rgef-1* using primers oCM59 and oCM159), creating BiFC expression vectors pCM34 and pCM36-37. Finally, *pat-4* and *unc-112* cDNAs were synthesized and provided in vectors pCM216 and pCM217 from IDT. The cDNA of *pat-4* was digested from pCM216 with BsiWI and cloned into BsiWI-digested pCM38 via T4 DNA ligase. The cDNA of *unc-112* was PCR amplified using primers oCM218-219 and cloned by Gibson Assembly into BsiWI-digested pCM39. For a list of all vectors, DNA fragments, and primers used in this study, please see **Table S2**.

**Figure 1:**
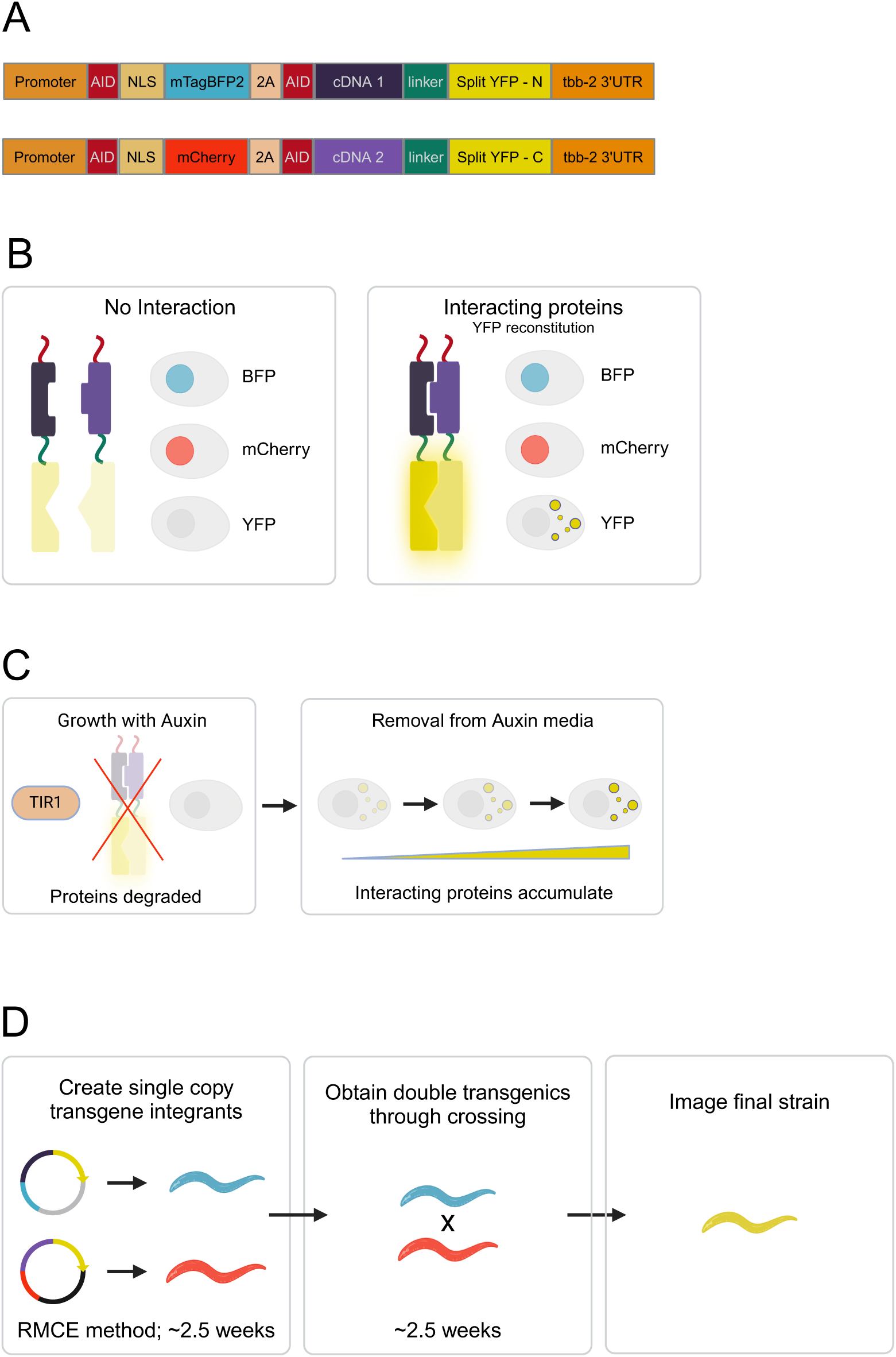
A revised and updated BiFC expression system for *C. elegans*. **A)** Diagram of both modified BiFC vectors. Each vector is expressed under a promoter of choice. A fluorescent reference protein, either mTagBFP2 or mCherry, is fused via 2A peptide to each half of the split-YFP protein. cDNAs encoding proteins of interest to be tested are inserted between the 2A peptide and YFP fragment, separated by a flexible linker. Additionally, the mTagBFP2 and mCherry proteins are tagged with a nuclear localization signal (NLS). Both fluorescent reference proteins and the protein of interest fused to the split-YFP are tagged with an auxin inducible degron. **B)** When each transgene is expressed and translated, the 2A peptide will self-cleave, creating two distinct fusion proteins. The mTagBFP2/mCherry will localize to the nucleus while the proteins of interest fused to the split-YFP half will localize to their endogenous subcellular location. If the two proteins of interest interact, the split-YFP halves will be brought together and reconstitute, resulting in split-YFP fluorescence. If the proteins do not interact there will be no split-YFP fluorescence. **C)** Introduction of the auxin inducible degron system allows for the conditional expression of the BiFC system. **D)** Schematic of rough timeline for BiFC assays once expression vectors have been cloned.

### DNA Preparation, microinjection, and RMCE single-copy transgenesis

Plasmids were prepared by a standard purification kit (GeneAid), followed by further purification using a clean and concentrator kit (Zymo) and eluted into water. L4 worms were picked 24 hours ahead of time and then adult worms were mounted on 2% agarose pad in a drop of halocarbon oil. Plasmids were injected using standard procedures at a final concentration of 50 ng/μL and diluted in water (Kadandale et al., 2009). After injection, animals were maintained and screened for single-copy insertions according to the RMCE protocol described in (Nonet, 2020). Integrated strains were confirmed by visualizing fluorescence of mCherry and mTagBFP2.

### Fluorescence Microscopy

Fluorescence microscopy micrographs were taken on a Leica TCS SP8 confocal microscope. L4 stage worms were mounted on 2% agarose pads and then immobilized with 5-10 mM levamisole diluted in water or M9. Animals were then imaged at 40X or 63X magnification using sequential illumination and scanning of fluorescent proteins. Images from 10-15 worms were collected for each sample and for further quantification.

### Quantification of fluorescence

All microscopy settings (laser power, gain, emission spectra) were kept the same in a given experiment comparing fluorescence intensity between strains. Settings were chosen to maximize the dynamic range of signal while also preventing saturation of pixels. Care was taken to ensure that only nuclei or cells without saturated pixels were quantified. Images were quantified using FIJI ImageJ software (Schindelin et al., 2012). First, a maximum intensity projection was created. For bFOS/bJUN measurements under the *rgef-1* promoter, neuronal nuclei of the ventral nerve cord near the vulva were measured. Therefore, only the slices that captured the ventral nerve cord were included in the maximum intensity projection. For bFOS/bJUN measurements under the *myo-3* promoter all slices were included in the maximum intensity projection. For bFOS/bJUN measurements, a circular region of interest (ROI) was drawn around each nucleus being measured and an equally sized ROI was measured beside each nucleus as a background measurement. Fluorescence was measured as an integrated density value. 10 nuclei and their respective control ROI were measured per animal. Corrected total cell fluorescence (CTCF) was calculated by the following equation: CTCF = Integrated Density_nucleus ROI_ – (Area_ROI_ * Mean Fluorescence_control ROI_).

### Auxin Inducible Degron Experiments

Standard NGM plates with an addition of auxin (Indole-3-acetic acid; Sigma-Aldrich). dissolved in ethanol to a final concentration of 25 μM were made and seeded with OP50-1 bacteria. Two L4s were plated per strain. Four days later around 60 F1 progeny at L3-L4 stages were moved from the auxin containing plates to standard NGM plates to restore accumulation of BiFC transgene proteins. Starting from this 0 hour time point, the fluorescence of 2-5 animals per strain was measured and quantified every 2 hours for a total of 12 hours using the fluorescence microscopy and quantification methods described above.

### Statistical Analysis

For pairwise-comparisons throughout the figures, the p-values were calculated using the two-sided Wilcoxon rank-sum test. No multiple hypothesis correction was used for most comparisons due to small number of comparisons made per experimental group. However, for Figure 5B and C, we report an adjusted p-value calculated using the two-sided Wilcoxon rank-sum test and a Bonferroni adjustment. Each scatter plot contains a smooth local regression line fitted to the data with a confidence interval of 0.95 shown.

## Results

### An improved and more flexible BiFC assay in C. elegans

Given recent advances in transgenesis approaches and in conditional expression tools in *C. elegans* (Ashley et al., 2021; Nonet, 2020; Zhang et al., 2015), we modified the original BiFC expression vector design to take advantage of these new features (**Fig. 1**). Our modified assay employs two vectors. One vector contains the mCherry red fluorescent protein tagged with a nuclear localization signal (NLS) and the C-terminal half of Venus (VC155), a yellow fluorescent protein (YFP) (**Fig. 1A**). The second vector contains the mTagBFP2 blue fluorescent protein tagged with a NLS and the N-terminal half of Venus (VN177) (**Fig. 1A**). In each expression vector, the additional fluorescent proteins and split YFP fragments are separated by a 2A peptide (**Fig. 1A**). The 2A peptide allows both proteins to be translated in the same reading frame but cleaved into two separate proteins after the ribosome translates through the 2A peptide sequence (Ahier & Jarriault, 2014; Kim et al., 2011). The mCherry and mTagBFP2 proteins serve as controls to label cells expressing the split-YFP transgenes and also to normalize split-YFP signals across individual cells (**Fig. 1B**).

Upstream of the split YFP coding sequence is a multiple cloning site which allows for the insertion of cDNAs encoding the proteins of interest (**Fig. 1A**). Currently the vectors are designed to tag the C-terminus of the query proteins with one half of the split-YFP, but this fusion is bridged by a flexible linker to minimize the impact of the fluorescent protein tag on the function of each protein and their potential interaction (Hiatt et al., 2008; Shyu et al., 2008).We also fused auxin inducible degradation (AID) tags (degrons; (Zhang et al., 2015)) to the mCherry, mTagBFP2 and both halves of the cDNA of interest::split-YFP fusions (**Fig. 1A**). The addition of the AID degrons allow for auxin-controllable degradation of all fluorescent proteins, enabling temporal control of the split-YFP reconstitution (**Fig. 1C**). Finally, another cloning site is available at the 5′ end of the expression cassette to insert promoters of interest (**Fig. 1A**). To date, we have designed vectors that drive expression broadly in the nervous system (*rgef-1* promoter) or in body wall muscle cells (*myo-3* promoter).

The original BiFC vectors are designed for multi-copy expression in *C. elegans* and maintained through extrachromosomal arrays (Hiatt et al., 2008; Shyu et al., 2008). However, the overexpression and variability of extrachromosomal arrays make it difficult to compare signal between animals (both same strain and controls) and leads to high levels of baseline YFP-reconstitution, even in the absence of additional fused proteins. Moreover, over-expression of query proteins of interest can lead to artifacts such as aggregation or mis-localization. To address some of these caveats, we have inserted our BiFC expression cassettes into the pLF3FShc vector, which enables single copy integration of our transgenes into the genome using FLP Recombinase Mediated Cassette Exchange (RMCE) (Nonet, 2020). Several RMCE recipient strains are available, with FRT recombination sites located on different chromosomes, facilitating the integration of both of our transgenes at distinct genomic locations. Using the RMCE integration method followed by genetic crosses to place both transgenes in the same strain, imaging experiments can be performed as early as 5 weeks after vectors are created (**Fig. 1D**).

In the sections that follow, we describe our experiments testing the additional features included in our updated BiFC expression vectors.

### Quantifying protein interactions in single neurons and muscle cells with modified BiFC

We first tested our BiFC expression vectors with a well-characterized protein-protein interaction. Specifically, we used cDNAs encoding the basic region leucine zipper domains of mammalian FOS (bFOS) and JUN (bJUN), two transcription factors that are known to form a heterodimer in the nucleus through this domain (Hiatt et al., 2008; Shyu et al., 2008). Driving expression of bFOS and bJUN in neurons or body wall muscle cells, we observed clear nuclear mCherry, mTagBFP2, and split-YFP signal in individual cells (**Figs. 2A,B and 3A,B**). Previous studies demonstrated that the bFOS-bJUN PPI can be disrupted by deleting a leucine zipper domain of bFOS (bFOSΔZIP) (Hiatt et al., 2008). Consistent with these findings, despite strong mCherry and mTagBFP2 signal, we observed little to no split-YFP fluorescence in neurons and muscle cells when bJUN and the mutated bFOSΔZIP were co-expressed (**Figs. 2A,B and 3A,B**). Additionally, we did not detect any split-YFP fluorescence when our empty (i.e without any cDNAs of interest) BiFC reporters were expressed in neurons or muscle cells (**Fig. S1**). Taken together, our modified BiFC split-YFP transgenes faithfully report the subcellular location of a known PPI, and the strength of reconstituted signal is sensitive to alterations in interaction strength.

**Figure 2:**
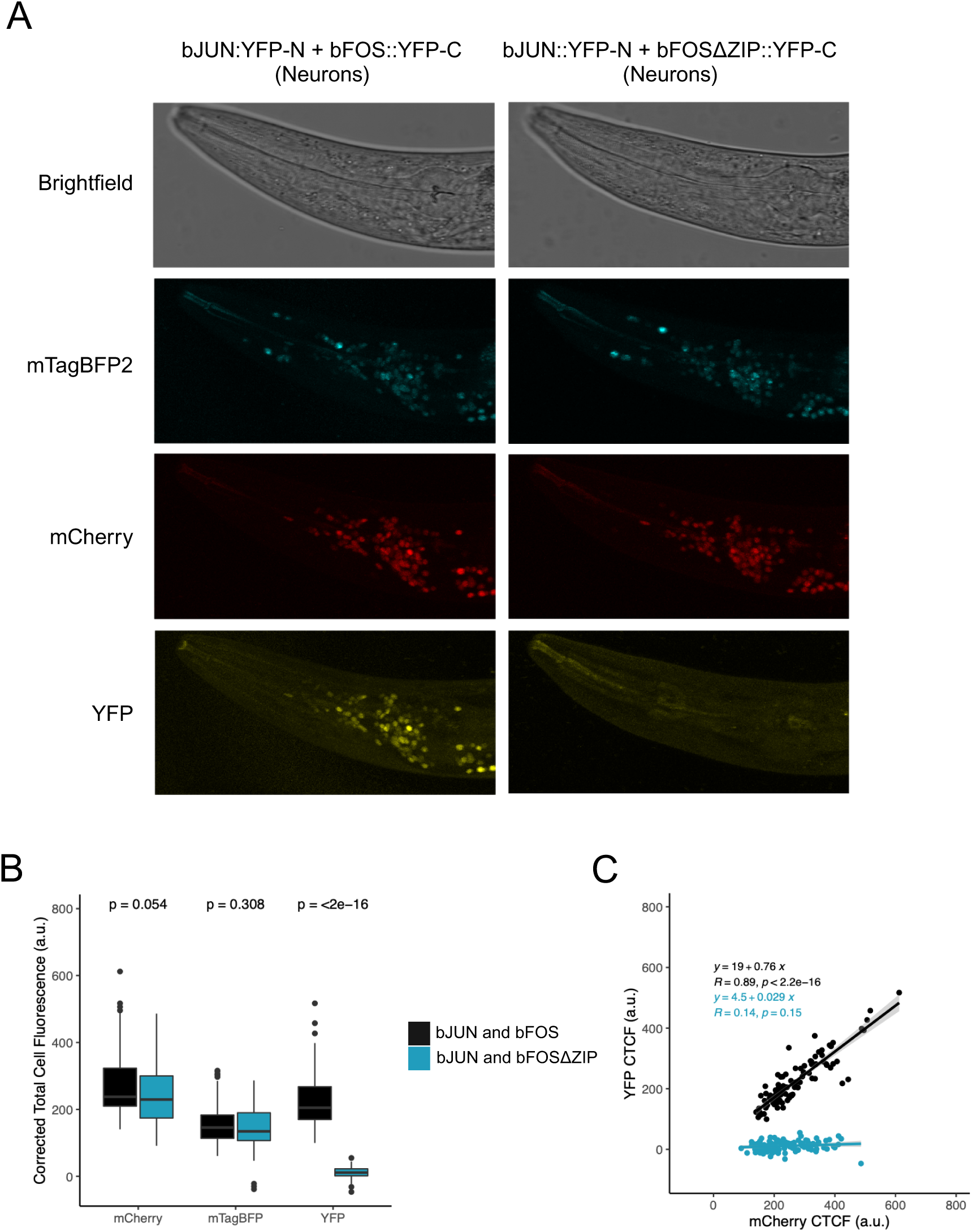
BiFC recapitulates the bJUN/bFOS protein interaction in neurons. **A)** Confocal fluorescent microscopy Z-stack micrographs showing brightfield, mTagBFP2, mCherry and YFP channels of the *rgef-1* promoter expression of mTagBFP2::2A::bJUN::YFP-N and mCherry::2A:: bFOS or bFOSΔZIP::YFP-C constructs. **B)** Boxplot of corrected total cell fluorescence (CTCF) of each fluorescent protein (mCherry, mTagBFP2 and split-YFP), comparing the CTCF of worms expressing bJUN and bFOS fused split YFP halves to the split-YFP fluorescence of bJUN and bFOSΔZIP fused split-YFP halves under the *rgef-1* promoter. P-values were calculated using the two-sided Wilcoxon rank-sum test. **C)** Scatter plot of the correlation between YFP fluorescence and the mCherry proteins for both the bJUN and bFOS positive and negative control. Contains a smooth local regression line with a confidence interval of 0.95. (n = 10 animals and 10 nuclei per animal for each condition)

**Figure 3:**
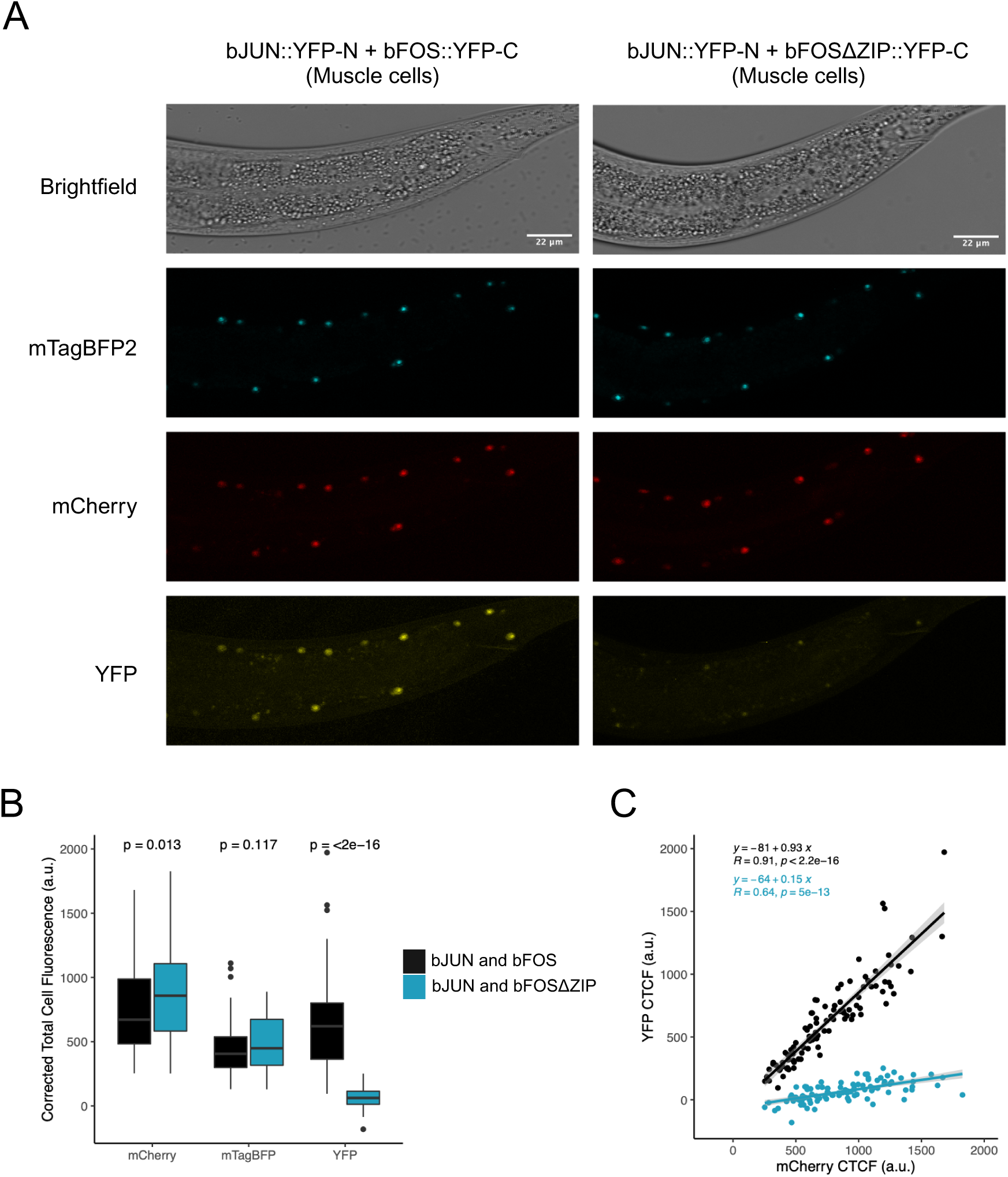
BiFC recapitulates the bJUN/bFOS protein interaction in muscle cells. **A)** Confocal fluorescent microscopy Z-stack micrographs showing brightfield, mTagBFP2, mCherry and YFP channels of the *myo-3* promoter expression of mTagBFP2::2A::bJUN::YFP-N and mCherry::2A::bFOS/bFOSΔZIP::YFP-C constructs. **B)** Boxplot of corrected total cell fluorescence (CTCF) of each fluorescent protein (mCherry, mTagBFP2 and split-YFP), comparing the CTCF of worms expressing bJUN and bFOS fused split YFP halves to the split-YFP fluorescence of bJUN and bFOSΔZIP fused split-YFP halves under the *myo-3* promoter. P-values were calculated using the two-sided Wilcoxon rank-sum test. **C)** Scatter plot of the correlation between YFP fluorescence and the mCherry proteins for both the bJUN and bFOS positive and negative control. Contains a smooth local regression line with a confidence interval of 0.95. (n = 10 animals and 10 nuclei per animal for each condition)

As stated above, our adoption of the RMCE transgenesis approach enabled our BiFC expression transgenes to be expressed at single copy, mitigating the variability in fluorescent signal between animals and even between individual cells associated with multi-copy mosaic over-expression from extrachromosomal arrays. As expected, we found that the overall distribution of fluorescence signals from the nuclear mCherry and mTagBFP2 marker proteins were not significantly different (neuronal expression; **Fig. 2B**) or only modestly different (muscle cell expression; **Fig. 3B**) when comparing promoter-matched strains expressing different combinations of bFOS/bJUN BiFC transgenes. However, even with single-copy integration of transgenes at defined loci, we found that the intensities of these reference fluorescent protein markers varied by nearly three-fold across individual cells, likely reflecting differential promoter activity (**Figs. 2C, 3C, and Fig. S2**).

Importantly, there was a positive linear correlation between the reconstituted bFOS/bJUN split-YFP and reference fluorescent protein signals (**Figs. 2C**, r = 0.89; **Fig. 3C**, r = 0.91). This strong correlation suggests that single cell split-YFP measurements can be normalized by reference fluorescent protein signals. Thus, we reasoned that the slope of the line of best fit through these data points can serve as a normalized measure of interaction signal within a tissue (**Figs. 2C and 3C**). Comparing the regression line slopes for the different bFOS/bJUN pair-wise interactions, we observed a 26.2 fold increased relative interaction signal of the bFOS/bJUN interaction compared with bFOSΔZIP/bJUN when expressed in neurons (slope_bFOS/bJUN_ / slope_bFOSΔZIP/bJUN_ = 0.76 / 0.029; **Fig. 2C**). In muscle cells, the bFOS/bJUN relative interaction signal was 6.2 fold higher compared with bFOSΔZIP/bJUN (slope_bFOS/bJUN_ / slope _bFOSΔZIP/bJUN_ = 0.93 / 0.15; **Fig. 3C**). Collectively, our results suggest that our single-copy expressed bicistronic BiFC transgenes enable the normalization of split-YFP signals across single cell measurements, leading to a more quantitative readout of protein-protein interactions within tissues of interest.

### Visualizing protein-protein interactions at endogenous subcellular foci

To confirm that our modified BiFC system has the ability to visualize the subcellular localization of an interaction in the cytoplasm, we inserted the cDNAs encoding the *C. elegans* proteins UNC-112 and PAT-4 into our muscle expression vectors. UNC-112 and PAT-4 both have integrin binding activity and interact with each other at the dense bodies and m-line of muscle fibers in *C. elegans* (Mackinnon et al., 2002; Qadota et al., 2012; Rogalski et al., 2000). As a negative control, we co-expressed UNC-112 and bFOS, which are not known to interact and localize to different subcellular compartments.

We observed nuclear mCherry and mTagBFP2 signal in the body wall muscles and reconstituted split-YFP fluorescence in dense bodies and m-lines in the worms co-expressing UNC-112 and PAT-4 BiFC transgenes (Francis & Waterston, 1985) (**Fig. 4 and Fig. S3**). In contrast, no visible reconstituted split-YFP fluorescence was observed in the negative control (**Fig. 4**). These results indicate that known protein-protein interactions in cytoplasmic subcellular organelles can also be detected by our BiFC reporters. Moreover, the partitioning of nuclear mCherry and mTagBFP2 signals from the cytoplasmic split-YFP signal (**Fig. 4**) suggests that the nuclear localization signals in our bicistronic reporters do not interfere with endogenous localization patterns of the proteins to be tested. This latter observation is consistent with results from the original study implementing 2A peptides in *C. elegans* (Ahier & Jarriault, 2014).

**Figure 4:**
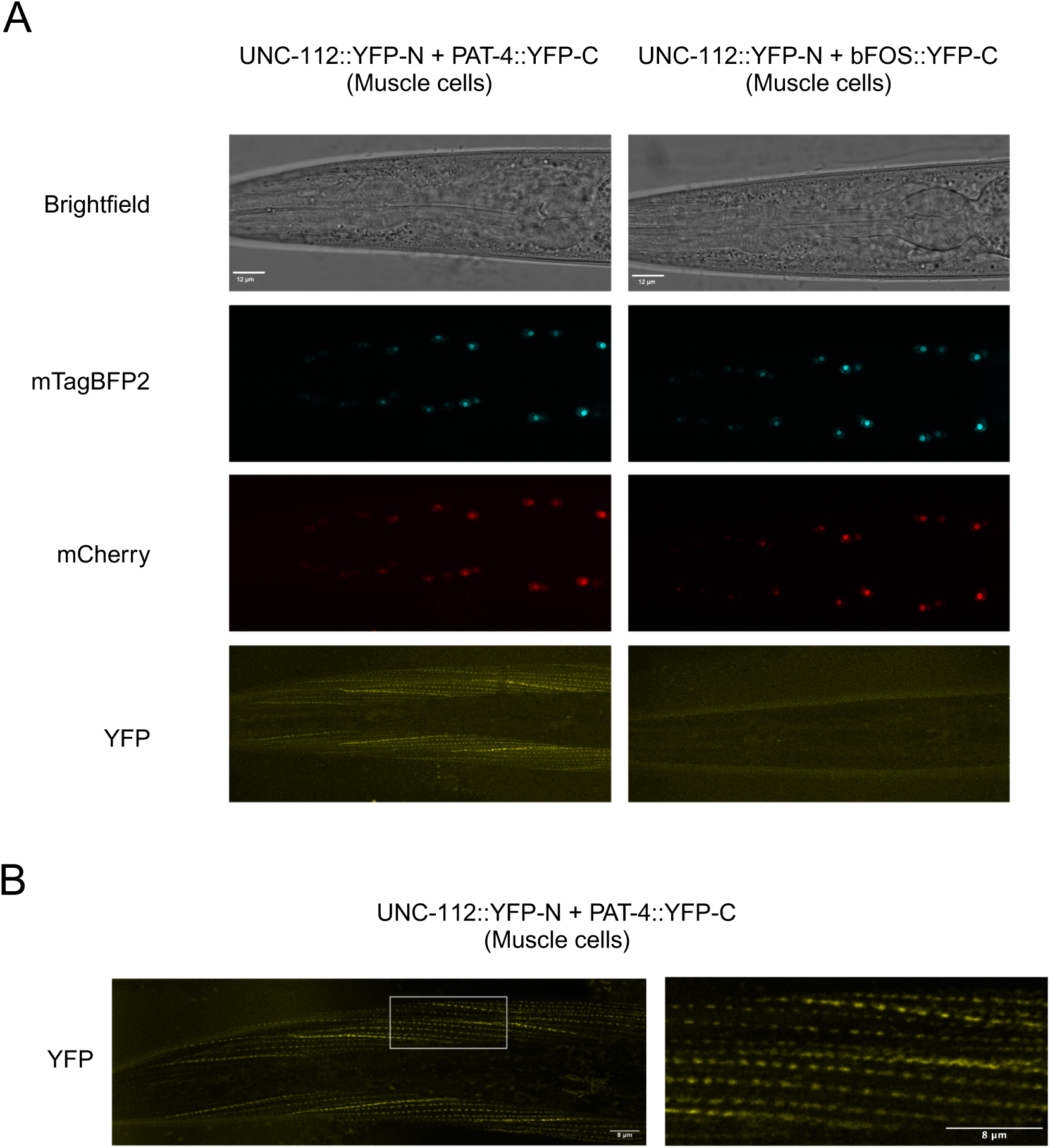
Visualization of a known subcellular endogenous interaction between PAT-4 and UNC-112 proteins at the body wall muscle dense bodies. **A)** Confocal fluorescent microscopy Z-stack micrographs showing brightfield, mTagBFP2, mCherry and YFP channels of the *myo-3* promoter expression of mTagBFP2::2A::UNC-112::YFP-N and mCherry::2A::PAT-4::YFP-C constructs and mTagBFP2::2A::UNC-112::YFP-N and mCherry::2A::bFOS::YFP-C constructs . **B)** Deconvolution image of confocal fluorescent microscopy Z-stack micrographs showing YFP channels of the *myo-3* promoter expression of mTagBFP2::2A::UNC-112::YFP-N and mCherry::2A::PAT-4::YFP-C constructs. Whitebox indicates magnified region visualized.

### Coupling BiFC with the auxin-inducible degron system enables spatio-temporal detection of protein interactions

As described above, a limitation to the BiFC approach is that reconstitution of the two halves of split-YFP are irreversible (Hiatt et al., 2008; Shyu et al., 2008). Thus, in some cases, robust fluorescent signals can be observed even in negative control reporters when measured at steady state (Hiatt et al., 2008; Shyu et al., 2008). To enable inducible control over the accumulation of reconstituted YFP signal, we have fused auxin inducible degradation (AID) tags (degrons) to mCherry, mTagBFP2 and both halves of the split YFP in our reporters (**Fig. 1**; (Zhang et al., 2015). This degron can be recognized by the plant E3 ubiquitin ligase TIR1 only in the presence of the small molecule auxin, allowing for the conditional depletion of proteins of interest (Zhang et al., 2015).

To test our inducible BiFC strategy, we crossed animals expressing the bJUN and bFOS/bFOSΔZIP controls under the *myo-3* promoter into a strain expressing TIR1 under the pan-somatic *eft-3* promoter (Ashley et al., 2021). Animals were grown on plates containing 25 uM of auxin for a generation, and their L3-L4 progeny were then transferred to standard growth plates lacking auxin (t = 0 hours) and imaged every 2 hours for 12 hours (**Fig. 5 and Fig. S4**). As expected, immediately after removal from auxin (t = 0 hours), we observed little to no fluorescence for any of the fluorescent proteins (**Fig. 5**). In as little as four hours post-auxin removal, we observed a significant increase in mCherry reference protein fluorescence in both the bFOS/bJUN and bFOSΔZIP/bJUN expressing strains, with levels continuing to rise over the 12 hour time course (**Fig. 5A and B, Fig. S5**). As expected, the split-YFP fluorescence returned in the bFOS/bJUN expressing animals, showing significant differences from baseline measurement (t = 0 hours) as early as 4 hours post-auxin removal, reaching a maximum signal at 12 hours (**Fig. 5A and C, Fig. S5**). In contrast, in bFOSΔZIP/bJUN expressing animals, split-YFP signal remained close to baseline levels throughout the time course (**Fig. 5A and C, Fig. S5**). Previous studies have found that some degradation of AID-tagged proteins can occur with co-expression of TIR1 in the absence of auxin (Negishi et al., 2022; Schiksnis et al., 2020). We found this was also the case in with our BiFC reporters, where split-YFP and reference fluorescent protein signals were significantly lower in TIR1-expressing strains compared with strains lacking TIR1 for both the positive and negative control (**Fig. S5**).

**Figure 5:**
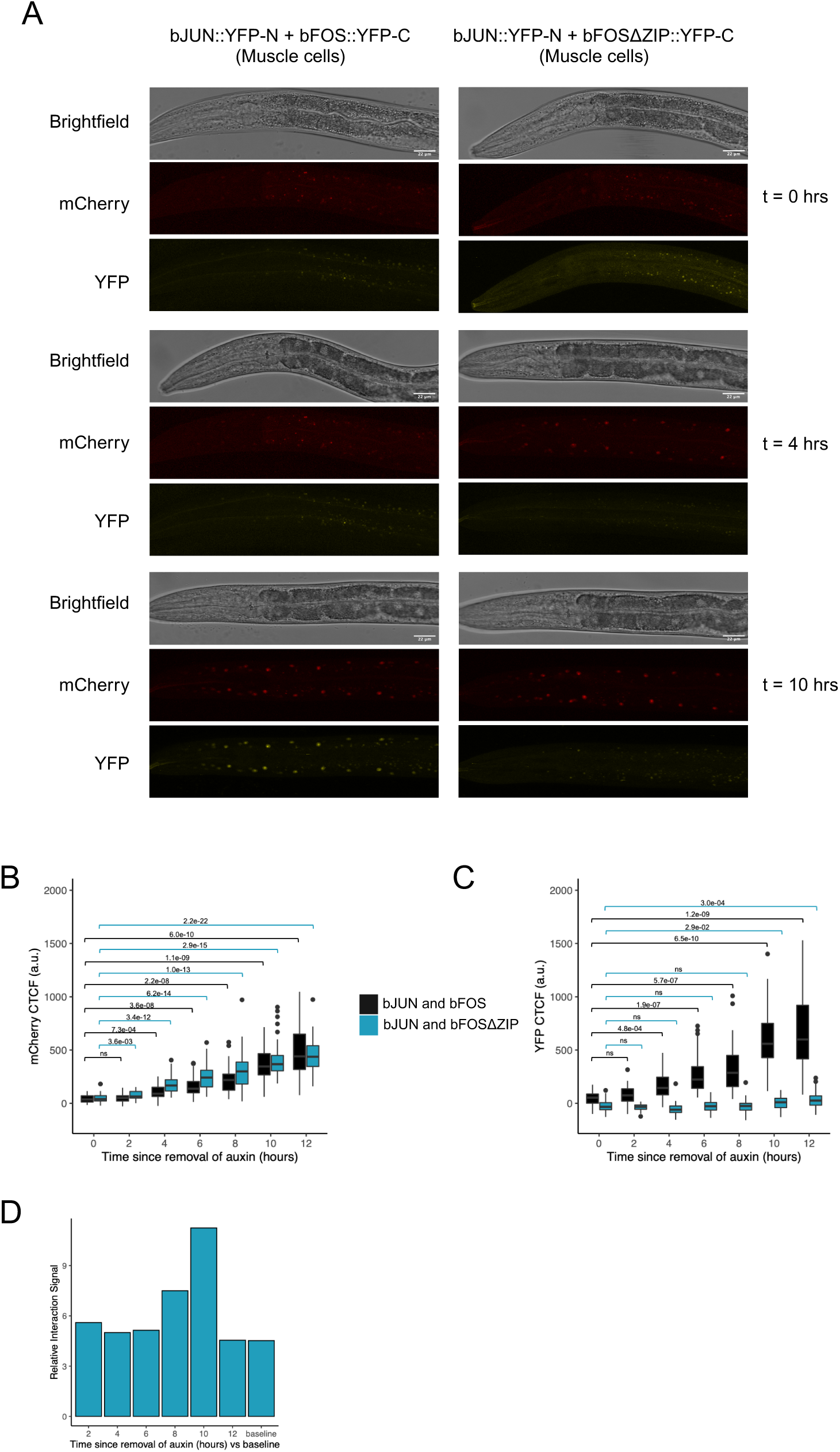
Adopting the AID system to monitor BiFC interactions in an inducible manner. **A)** Confocal fluorescent microscopy Z-stack micrographs showing brightfield, mCherry and YFP channels of the *myo-3* promoter expression of mTagBFP2::2A::bJUN::YFP-N and mCherry::2A::bFOS/bFOSΔZIP::YFP-C constructs co-expressing eft-3p::TIR1, 0, 4 and 10 hours after worms were removed from the presence of 25 uM of auxin. **B-C)** Boxplot showing the CTCF of the mCherry and split-YFP fluorescent proteins of worms expressing under the *myo-3* promoter bJUN and bFOS and bJUN and bFOSΔZIP fused to split-YFP halves and co-expressing the protein TIR1. The CTCF was measured every 2 hours after the worms were removed from the presence of 25 uM of auxin. (n= 2-5 animals and 10 nuclei per timepoint). Adjusted p-values were calculated using the two-sided Wilcoxon rank-sum test and a Bonferroni adjustment. **D)** Bar graph of the relative interaction signal (fold change) at each timepoint of the AID experiment and at steady state expression (baseline).

We next investigated if the relative interaction signal (slope_bFOS/bJUN_ / slope_bFOSΔZIP/bJUN_) differed over the time course and as the fluorescent signal returned to steady state levels after the removal of auxin (**Fig. 5D and Fig. S5**). Interestingly, the bFOS/bJUN relative interaction signal reached a maximum of 11.25 fold higher than that of bFOSΔZIP/bJUN at ten hours post-auxin removal (slope_bFOS/bJUN_ / slope _bFOSΔZIP/bJUN_ = 1.8 / 0.16; **Fig. 5D and Fig. S5).** This is a 2.5 fold increase compared to a steady-state relative interaction signal of 4.5 (slope_bFOS/bJUN_ / slope _bFOSΔZIP/bJUN_ = 1.9 / 0.42; **Fig. 5D and Fig. S5).** Notably, the relative interaction signal at twelve hours post-auxin removal decreases to a value of 4.5, similar to our steady state measurements (slope_bFOS/bJUN_ / slope_bFOSΔZIP/bJUN_ = 1.5 / 0.33; **Fig. 5D and Fig. S5).** Taken together, our results indicate that our BiFC reporters and protein interaction signals can be made inducible through pairing with existing TIR1-expressing reporter strains. Our data also suggests that background split-YFP fluorescence and steady-state signals can influence the normalized interaction signal measurements, highlighting the added value of collecting BiFC data over an inducible time course.

## Discussion

### A quantitative, inducible, fluorescence-based protein-protein interaction assay for C. elegans

Through our modifications to the original BiFC system, we have expanded the current toolbox for the study of PPIs in *C. elegans*. These modifications have created a PPI assay that is quantifiable and allows for the spatial and temporal control of an interaction of interest at its endogenous subcellular location. We anticipate that this improved system will allow future users to answer questions related to the characterization of PPIs that were previously difficult to study.

Our introduction of the relative interaction signal value provides a framework for quantification of a candidate PPI in different applications. For example, it is documented that distinct protein isoforms generated by alternative splicing can influence the strength of rather than a binary gain or loss of PPIs (Buljan et al., 2012; Corominas et al., 2014; Ellis et al., 2012; Yang et al., 2016). As such, different isoform combinations of known interacting partners could be tested for quantitative effects on interaction strength in our assay. Similarly, targeted mutagenesis experiments aimed at testing critical residues (such as post-translational modification sites (Wang et al., 2022) or even entire domains in endogenous tissues could be assessed for their impact on PPI strength. This greatly expands the original BiFC assay, which was previously used as a qualitative readout of PPIs.

The introduction of the auxin inducible degradation system into the BiFC assay enables temporal control of the split-YFP signal. From a technical perspective, controlled induction of fluorescence allows the experimenter the opportunity to observe the dynamics of split-YFP signal accumulation until steady-state levels are reached, optimizing interaction signal to noise values (**Fig. 5 and Fig. S5**). Perhaps more interestingly, this temporal control will also facilitate the characterization of dynamic PPIs at different developmental stages without concern about steady-state accumulation of signal from earlier time windows.

### A complementary fluorescence-based assay for testing PPIs: comparisons and considerations

We envision that our modified BiFC approach will serve a complementary role with other approaches that assay PPIs in *C. elegans*. In particular, our approach could be used for focused validation studies that would follow from large scale approaches such as affinity purification or proximity labeling (i.e TurboID) coupled with mass spectrometry (Branon et al., 2018; Remmelzwaal & Boxem, 2019). Similarly, PPIs identified by yeast two hybrid studies could also be verified *in vivo* and in physiologically relevant tissues with our BiFC approach.

Recently, an interesting light-inducible PPI assay has been developed for use in *C. elegans* called CeLINC (Kroll et al., 2021). Like our modified BiFC approach, CeLINC enables the detection of binary PPIs in endogenous *C. elegans* tissues. One key difference is that CeLINC can take advantage of the growing library of community-generated fluorescent protein-tagged endogenous genes. In our BiFC method, cDNAs of interest are cloned into our expression vectors and integrated into the genome using the RMCE method. Thus, if two proteins of interest are already tagged with different fluorescent proteins, CeLINC could be rapidly employed relative to our BiFC approach (Kroll et al., 2021). However, it should be noted that if different allelic variants need to be tested, new CeLINC assays would require additional transgenesis and/or genome editing of existing tagged loci.

Another key difference is that in CeLINC, the interaction is detected at oligomerized protein clusters that are not at the native subcellular location of the PPI, whereas with BiFC the PPI is detected *in situ*. Therefore, if it is suspected that a PPI requires or is influenced by the physiological compartment, it may be more desirable to use the BiFC assay. Additionally, the two methods allow for inducible detection of a PPI through different mechanisms. CeLINC uses light-activated oligomerization, which can allow detection within minutes post-induction (Kroll et al., 2021). In comparison, our BiFC approach uses the auxin inducible degron approach, which enables detectable PPI signal within a few hours post-induction. Taking these induction methods and times into consideration should guide the experimenter on a suitable method to meet the needs of the experiment. Finally, to assay quantitative effects on the strength of a PPI, our BiFC approach and the development of the normalized interaction strength value should prove to be useful.

One general consideration worth noting in BiFC is that the signal observed is a result of the PPI *in situ*. However, when studying the impact of different protein variants (e.g. different alleles or splice variants) on PPIs, any observed changes in split-YFP signal could be due to indirect factors such as altered protein abundance, mis-folding or mis-localization among the variants tested. These possibilities should be considered when interpreting BiFC results. Additionally, since BiFC transgenes are tagged with a degron and split-YFP fragment, it is important to also consider whether the tag may have an impact on normal protein function and localization.

### Future prospects: expanding the color palette of BiFC

Protein-fragment complementation assays (PCAs) have emerged as a common strategy for studying binary PPIs (Petschnigg et al., 2011). PCAs involve fusing interacting proteins of interest to two halves of a split-protein which, when reconstituted, leads to a measurable response (Petschnigg et al., 2011). Our BiFC method is one such assay in which two proteins of interest are each tagged with N- and C-terminal portions of split-YFP and their interaction leads to a visual readout through the reconstitution of the fluorescent protein. Recently, there have been developments in the use of additional split-fluorescent GFP and worm-optimized mScarlet proteins, which have primarily been used to study protein expression and localization patterns in *C. elegans* (Cabantous et al., 2005; Calarco et al., 2025; Cuentas-Condori et al., 2023; Goudeau et al., 2021; He, Siwei et al., 2019; Hefel & Smolikove, 2019; Noma et al., 2017). Moreover, additional orthogonal split fluorescent proteins continue to be engineered (Hu & Kerppola, 2003; Miller et al., 2015).

We envision that any of these additional split fluorescent proteins could be adopted within the framework that we have created in our BiFC approach. This allows for the ability to tailor the BiFC method for situations when it could be disadvantageous to use split-YFP (for example, when using existing fluorescent subcellular markers). Additionally, co-expressing spectrally separable split fluorescent proteins could in principle allow multiple interactions to be monitored simultaneously, which would make it possible to study multiple components in a protein complex, or even the influence of one protein complex on another. This latter idea would further expand the utility of our approach.

## Data availability

Strains and plasmids are available upon request. Table S1 contains the genotypes of all strains used in this study. Table S2 contains the sequences of all primers, DNA fragments, and vectors used and generated in this study. All raw microscopy data and quantification is available upon request.

## Acknowledgments and Funding

We thank members of the Calarco Lab for detailed discussions surrounding the manuscript. This work was supported by an NSERC Discovery Grant and Canadian Institutes of Health Research (CIHR) Project Grants to J.A.C. Some strains were provided by the CGC, which is funded by NIH Office of Research Infrastructure Programs (P40 OD010440).

---

**Figure S1:**
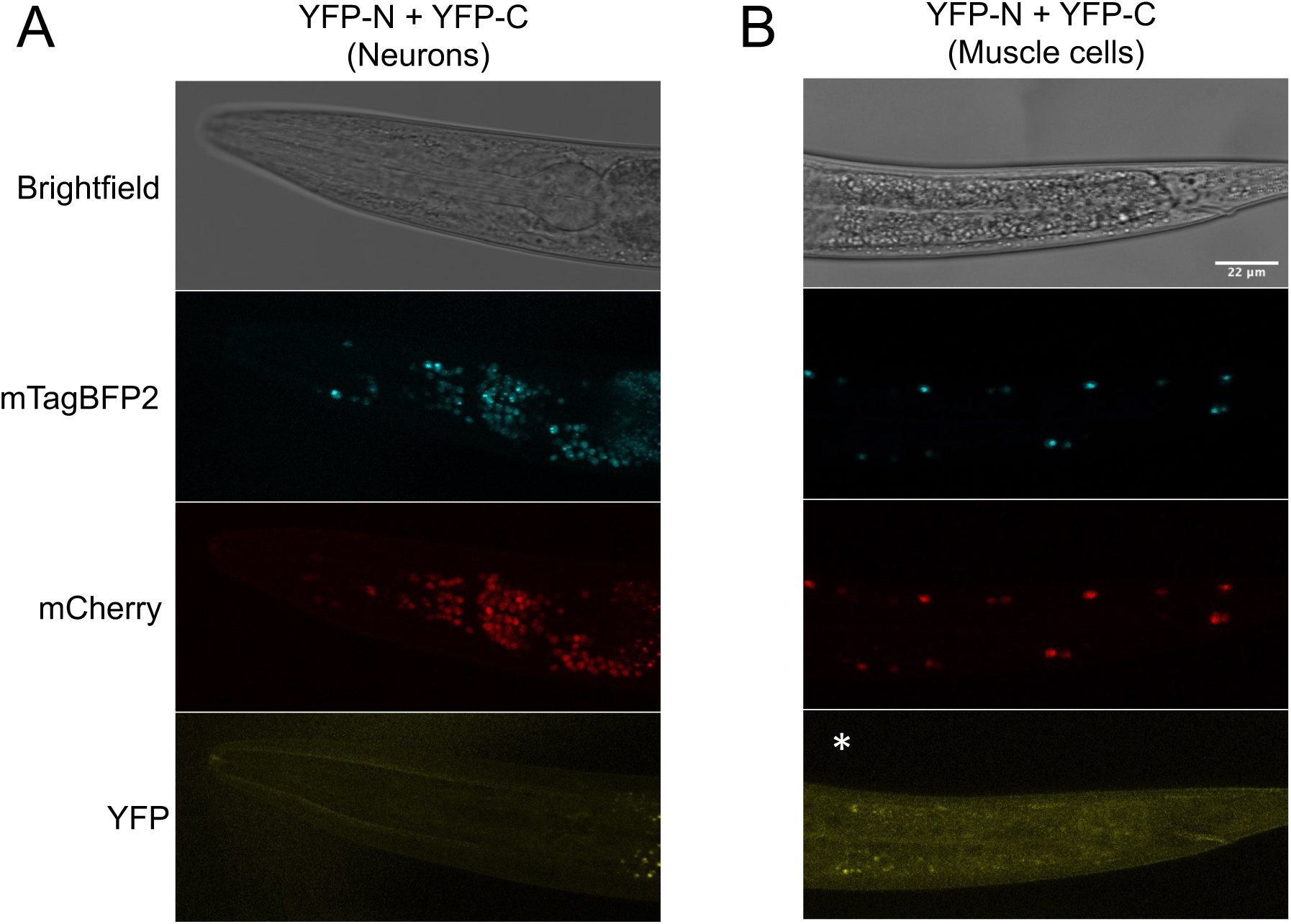
No split-YFP signal is visualized when the BiFC vectors are expressed fused to no cDNA. **A)** Z-stack maximum intensity projection micrographs showing brightfield, mTagBFP2, mCherry and YFP channels of the *rgef-1* promoter expression of mTagBFP2::2A::YFP-N and mCherry::2A::YFP-C constructs. **B)** Confocal fluorescent microscopy Z-stack micrographs showing brightfield, mTagBFP2, mCherry and YFP channels of the *myo-3* promoter expression of mTagBFP2::2A::YFP-N and mCherry::2A::YFP-C constructs. Asterisk denotes increased background fluorescence occasionally observed at periphery of Z-stacks when generating maximum intensity projections.

**Figure S2:**
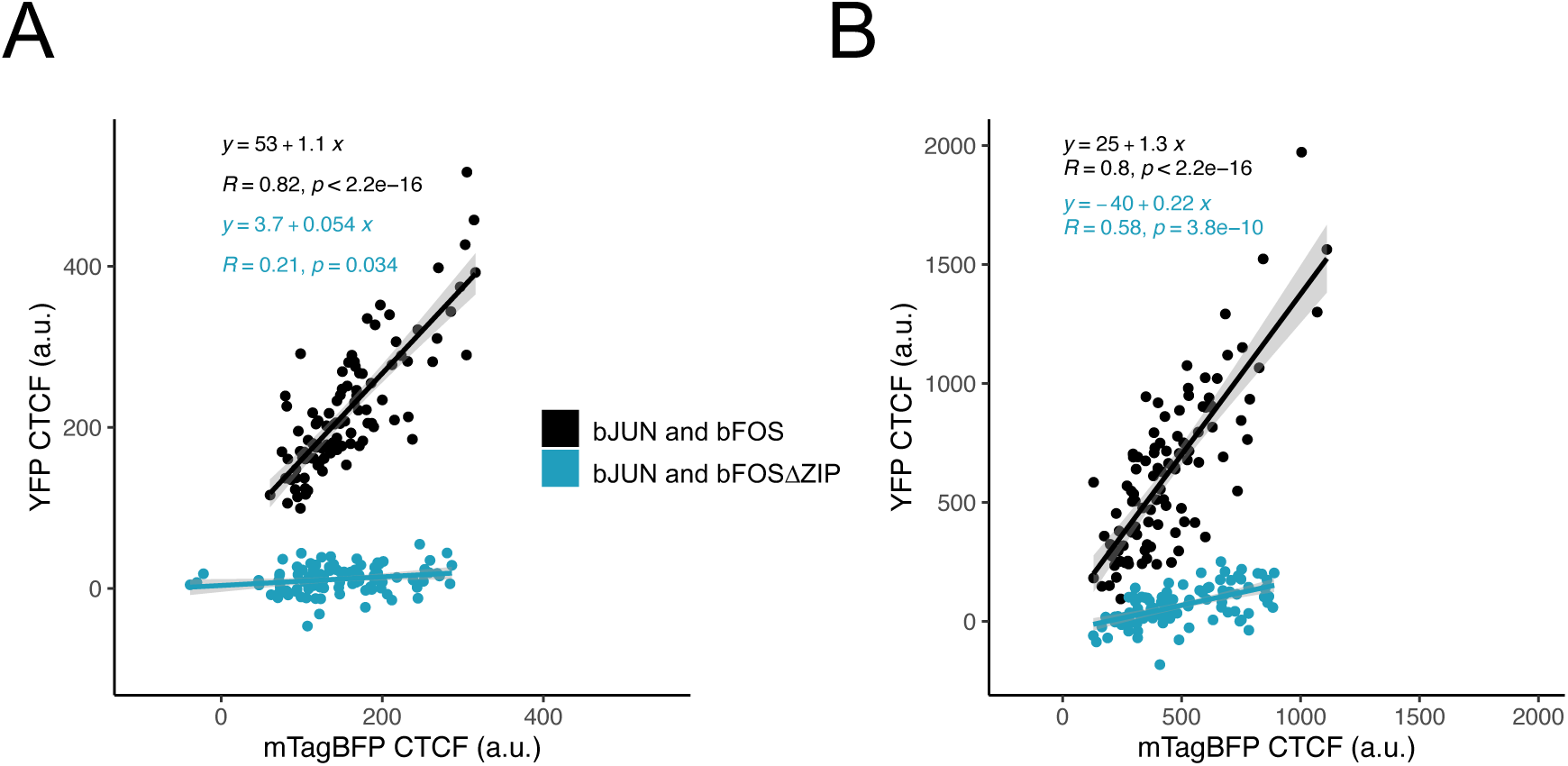
Correlation between YFP fluorescence and the mTagBFP2 proteins for both the bJUN and bFOS positive and negative control. **A,B)** Scatter plot of the correlation between *rgef-1* (**A**) *myo-3* (**B**) promoter expression YFP fluorescence and the mTagBFP2 proteins for both the bJUN and bFOS/bFOSΔZIP positive and negative control. Contains a smooth local regression line with a confidence interval of 0.95. (n = 10 animals and 10 nuclei per animal for each condition)

**Figure S3:**
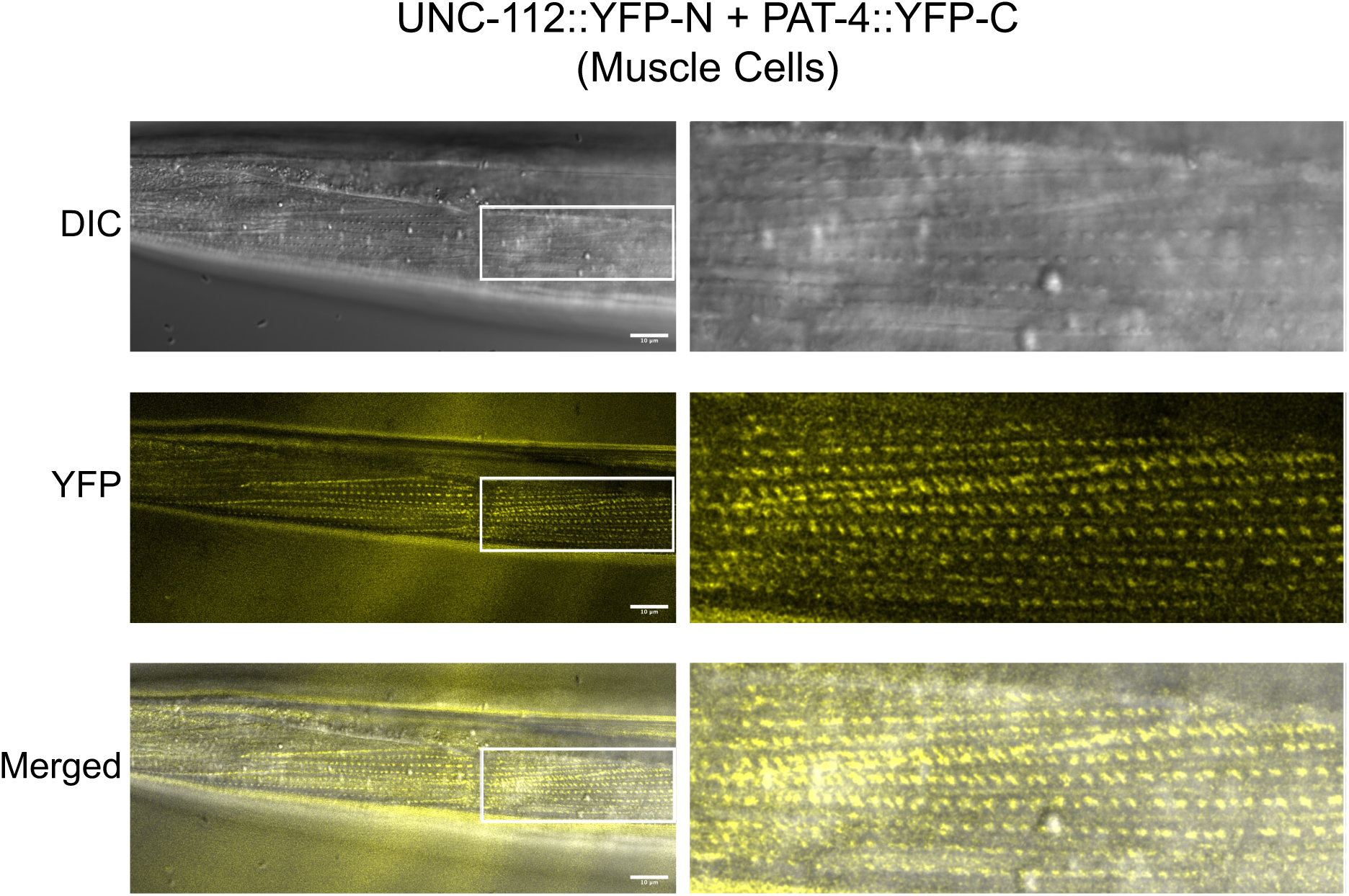
Observation of UNC-112 / PAT-4 interaction at dense bodies and M-lines. Deconvolution image of confocal fluorescent microscopy Z-stack micrographs showing DIC, YFP and merged channels of the *myo-3* promoter expression of mTagBFP2::2A::UNC-112::YFP-N and mCherry::2A::PAT-4::YFP-C constructs. Whitebox indicates magnified region visualized as presented on right panels.

**Figure S4:**
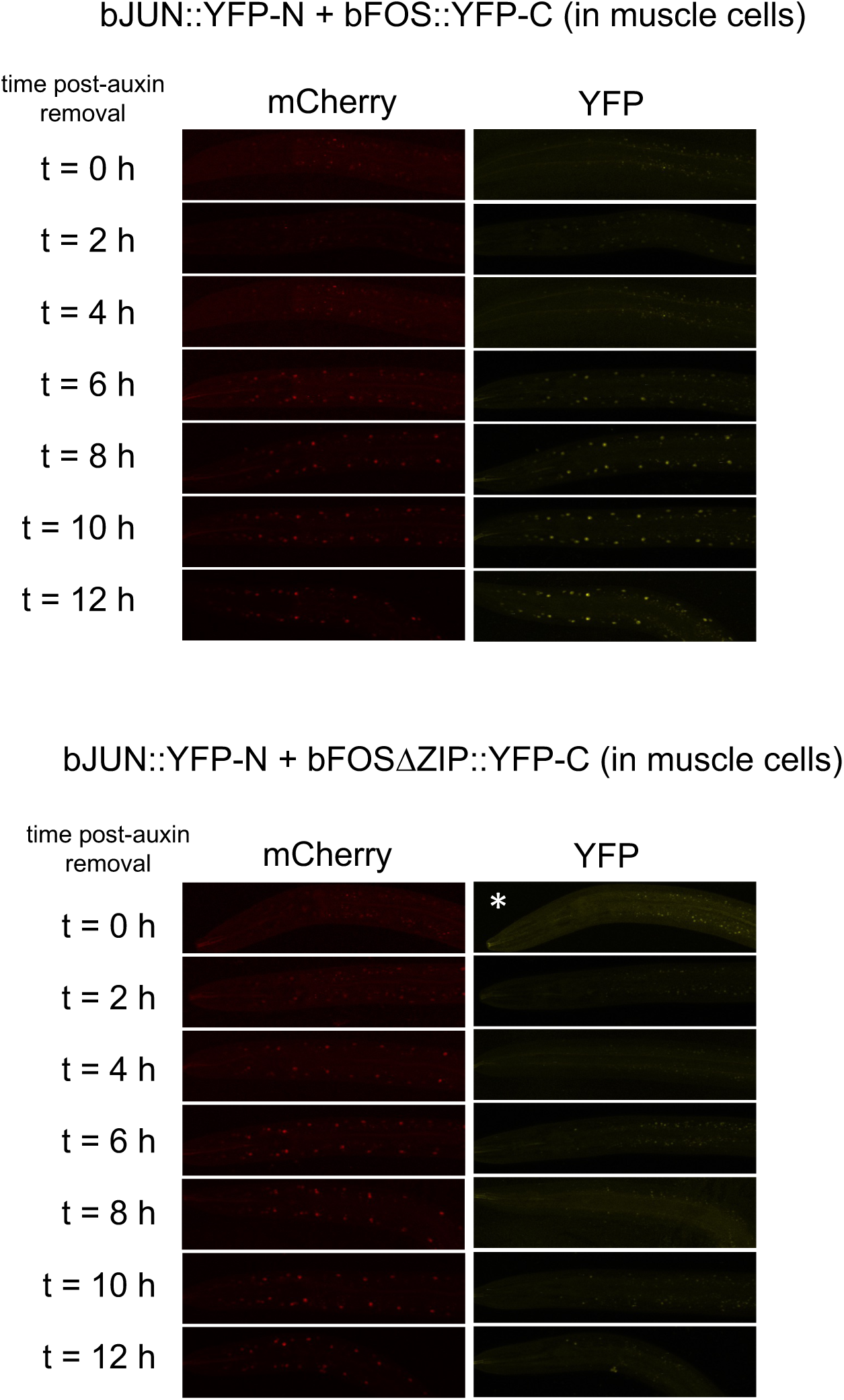
Full time course of AID experiment of inducible BiFC interactions. Z-stack maximum intensity projection micrographs showing mCherry and YFP channels of the *myo-3* promoter expression of mTagBFP2::2A::bJUN::YFP-N and mCherry::2A::bFOS/ bFOSΔZIP::YFP-C constructs co-expressing eft-3p::TIR1. Timepoints represent 0, 2, 4, 6, 8, 10 and 12 hours after worms were removed from the presence of 25 μM of auxin. Asterisk denotes increased background fluorescence occasionally observed at periphery of Z-stacks when generating maximum intensity projections.

**Figure S5:**
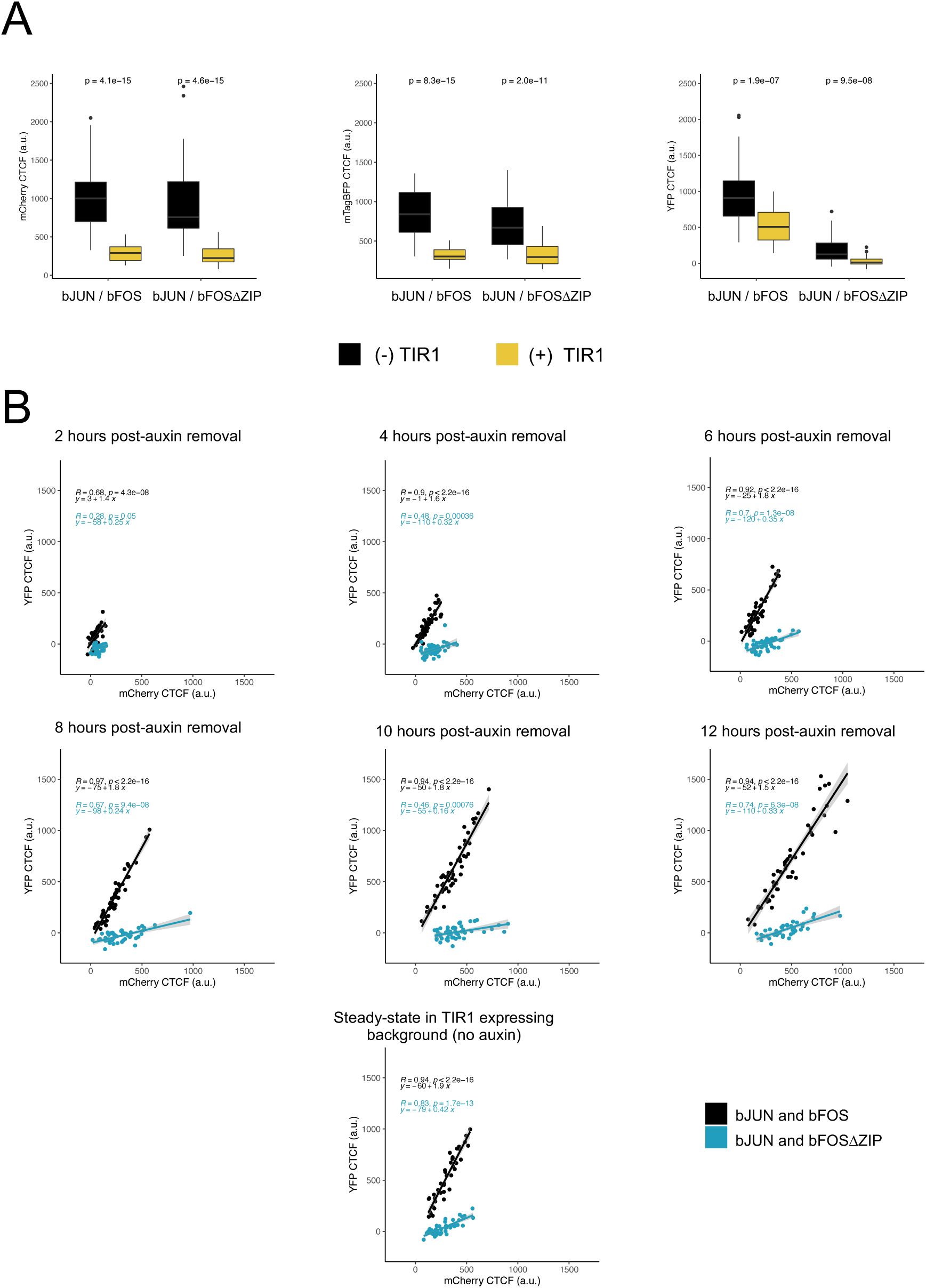
Influence of TIR1 on transgene expression and BiFC system fluorescent signal accumulation time course upon auxin removal. **A)** Boxplot of CTCF of each fluorescent protein (mCherry, mTagBFP2 and split-YFP), comparing the CTCF of worms expressing under the *myo-3* promoter either bJUN and bFOS or bJUN and bFOSΔZIP fused to the split YFP with or without the co-expression of eft-3p::TIR1, not in the presence of auxin. P-values were calculated using the two-sided Wilcoxon rank-sum test. (n = 4-5 animals and 10 nuclei per animal). **B)** Scatter plot of the correlation between YFP fluorescence and the mCherry proteins for both the bJUN and bFOS positive and negative control for each time point of the AID experiment and at steady state expression. Contains a smooth local regression line with a confidence interval of 0.95. (n = 4-5 animals and 10 nuclei per timepoint).

